# Both systemic Calcitonin Gene Related Peptide (CGRP) and a vestibular challenge promote anxiety-related behaviors and dynamic imbalance in mice

**DOI:** 10.1101/2023.06.30.547257

**Authors:** Shafaqat M. Rahman, Catherine Hauser, Stefanie Faucher, Elana Fine, Anne E. Luebke

## Abstract

Motion-induced anxiety and agoraphobia are more frequent symptoms in patients with vestibular migraine than migraine without vertigo. The neuropeptide calcitonin gene-related peptide (CGRP) is a therapeutic target for migraine and vestibular migraine, but the link between motion hypersensitivity, anxiety, and CGRP is relatively unexplored, especially in preclinical mouse models. To further examine this link, we tested the effects of systemic CGRP and off-vertical axis rotation (OVAR) on elevated plus maze (EPM) and rotarod performance in male and female C57BL/6J mice. Rotarod ability was assessed using two different dowel diameters: mouse dowel (r = 1.5 cm) versus rat dowel (r = 3.5 cm). EPM results indicate CGRP increased anxiety indexes and time spent in the closed arms in females but not males, while OVAR increased anxiety indexes and time spent in the closed arms in both sexes. The combination of CGRP and OVAR elicited even greater anxiety-like behavior. On the rotarod, CGRP reduced performance in both sexes on a mouse dowel but had no effect on a rat dowel, whereas OVAR had a significant effect on the rat dowel. Rotarod performance is influenced by dowel diameter, with larger dowels presenting greater challenges on balance function. These results suggest that both CGRP and vestibular stimulation induce anxiety-like behavior and that CGRP affects dynamic balance function in mice depending on the type of challenge presented. Findings highlight the potential translation of anti-CGRP receptor signaling therapeutics for treating motion hypersensitivity and motion-induced anxiety that manifests in vestibular migraine.

**Significance statement:** Anxiety is very common in patients with dizziness and vestibular migraine (VM). Elevated CGRP levels have been linked to migraine symptoms of increased light and touch sensitivity in mice and humans and we wondered if a systemic injection of CGRP into mice would increase anxiety and imbalance; and if mice further exposed to a vestibular stimulus would have their anxiety measures sharpened. We observed a female preponderance in both CGRP and motion-induced anxiety-like behaviors, suggesting that the role of CGRP in migraine’s anxiety symptoms can be recapitulated in the mouse. Our findings suggest that CGRP signaling has a pertinent role in motion-induced anxiety and dynamic imbalance, and warrants the potential use of anti-CGRP therapies for the treatment of these symptoms.

## Introduction

Anxiety is a common burden that arises with persisting dizziness in patients with vestibular disorders (Balci and Akdal 2020; Bigelow et al. 2016; Huang, Wang, and Kheradmand 2020). Patients with vestibular migraine (VM) have been reported to exhibit severe anxiety and neuroticism towards their condition based on outcomes from the Dizziness Handicap Inventory, Beck depression and anxiety scales, and Short Form health surveys (Ak et al. 2022). Elevated levels of anxiety are also associated with a 53% increased likelihood of falls based on a meta-analysis of eighteen studies linking anxiety as a risk factor for falls (Hallford et al. 2017). Fear of falling is largely detrimental to balance performance in older adults and vestibular patients.

The neuropeptide Calcitonin Gene-Related Peptide (CGRP) is involved in migraine and VM and is known to cause pain, various hypersensitivities, and anxiety (Hoskin and Fife 2021; Russo and Hay 2022). Recent studies have shown CGRP to induce light-aversion, tactile sensitivity, and nociceptive squinting behaviors (Wang et al. 2022; Wang et al. 2021). However, motion sensitivity due to CGRP is relatively unexplored in mouse models, even though motion-induced anxiety and agoraphobia – a fear of one’s surroundings – are more frequent symptoms in patients with vestibular migraine than migraine without vertigo (Kutay et al. 2017). Thus, this paper aims to study sex-specific changes in motion-induced anxiety and balance behavior in the C57BL/6J mice that are treated with systemically delivered CGRP and a challenging vestibular stimulus.

We used off-vertical axis rotation (OVAR) as the vestibular stimulus, since OVAR can be disorienting and can promote motion sickness and nausea in rodents and human healthy controls (Furman, Schor, and Schumann 1992; Idoux, Tagliabue, and Beraneck 2018; Zhang et al. 2022).

The elevated plus maze is a behavioral test extensively used to assess anxiety-like behavior in rodent models (Pellow et al. 1985). Measurements of the time spent in open versus closed arms and the number of entries (bouts) into each arm can be used to calculate an anxiety index, an indication of anxiety-like behavior that increases with a mouse’s greater presence in the closed arms (Cohen, Matar, and Joseph 2013). This test is used in preclinical research to evaluate anxiety-like behaviors caused by anxiogenic drugs like the known headache-inducing agent sodium nitroprusside (Al-Humaidhie, Thiab, and Hassen 2021).

The rotarod assay is commonly used to assess a mouse’s gait and balance on a rotating rod, and the mouse’s time (latency) to fall off the rod can provide behavior insights attributed to central and peripheral neurological dysfunction (Deacon 2013). Mice that lack αCGRP have been shown to perform poorly on rotarod (Jones et al. 2018).

To date, there are limited studies that have assessed CGRP-specific changes in mouse behavior that relies on vestibular cues (Zhang et al. 2020), and if CGRP-induced anxiety is exacerbated by vestibular provocation. Thus, this study’s objectives are to examine the role of intraperitoneally delivered CGRP and a vestibular stimulus in anxiety and dynamic balance behavior in the wildtype C57BL/6J mouse.

## Methods

### Animals

C57BL/6J mice were obtained from Jackson Laboratories (JAX 664) and Mice were housed under a 12 to 12 day/night cycle at the University of Rochester’s Vivarium under the care of the University of Rochester’s Veterinary Services personnel. They were kept in cages of up to 5 mice per cage and had ad libitum access to food, water, and bedding material. Equal numbers of male (M) and female (F) mice were tested, with a total of 76 mice (38M/38F) used in these studies. All animal procedures were approved by the University of Rochester’s IACUC committee and performed in accordance with NIH standards.

The studies were designed to have sufficient power to detect male/female differences. Before beginning the experiments each day, mice were acclimated to the testing room, which was maintained at a temperature of 22-23 degrees Celsius, for at least 30 minutes. They remained in this room until the experiments were completed. Injections were given after the acclimation period. Mice were tested at ages between 2.3 and 6 months for both the elevated plus maze and rotarod tests.

### Injections

Injections were given intraperitoneally (IP) using a 33-gauge insulin syringe and behavioral testing was carried out ∼20 minutes post-injection. PBS was used as the diluent and control injection. Rat α-Calcitonin Gene-Related Peptide (CGRP) (Sigma) was injected at a dose of 0.1 mg/kg.

### Elevated Plus Maze (EPM)

EPM measures anxiety by utilizing the inherent conflict between the exploration of a novel area and avoidance of its aversive features. The mouse is placed in the middle of the maze, which is an open square (5 × 5 cm) in the center of four arms (30 cm long x 5 cm wide with 2 enclosed and 2 open). The maze consists of opaque plexiglass panels and is mounted on a stand 30 cm high. A video camera records for 30 wutes of the mouse exploring the maze in regular, room lighting conditions. Mice have four five-minute sessions collected in two days. Mice were injected with either vehicle control (day 1) or CGRP (day 3) twenty minutes prior to the first EPM test. The mice would then be subjected to the vestibular challenge (see <u>Vestibular Challenge – Off-Vertical Axis Rotation (OVAR)</u> for methods) and would then be tested again for EPM activity. The session is scored to determine the animal’s tendency for exploring enclosed versus open arms, and the time spent in each area and the number of bouts into each arm are recorded. An anxiety index per animal is computed per experimental condition (**Fig. 1A**). Following the recording, the mouse is returned to its home cage. The maze is cleaned between subjects using an animal disinfectant (Rescue).

**Figure 1:**
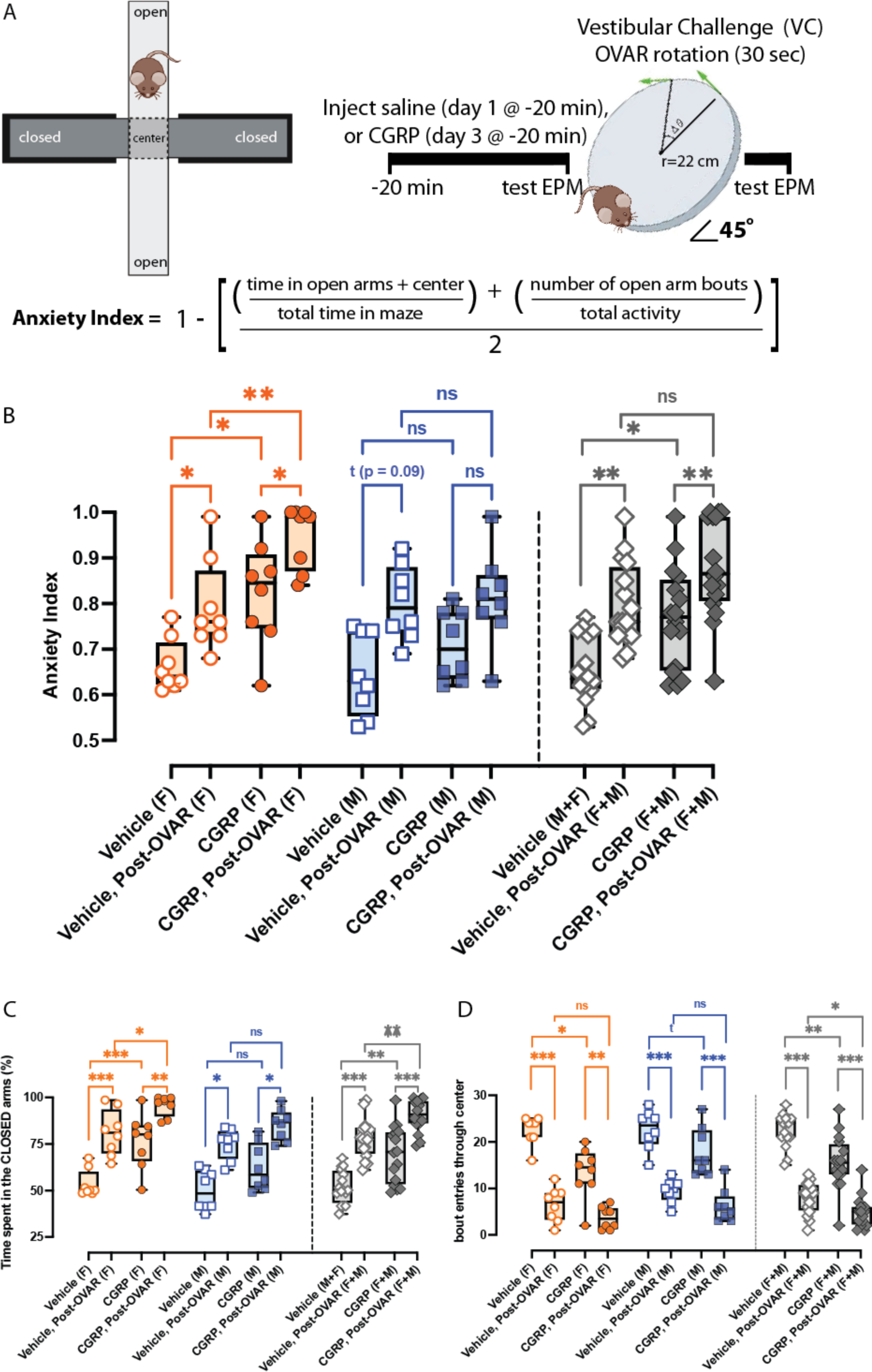
Calcitonin-Gene Related peptide (CGRP) and off-vertical axis rotation (OVAR) on elevated plus maze (EPM) behaviors. A, The EPM and its components are shown on the left, an experimental timeline is provided on the right, and an equation for calculating the anxiety index is provided underneath. **B-D**, In 16 mice (8M/8F), two-way repeated measures (RM) ANOVAs with Tukey post hoc analyses were used to evaluate **B**, anxiety indexes, **C**, time in closed arms, and **D**, center bouts due IP CGRP a brief off-vertical axis rotation (OVAR). Symbols are depicted: female (F) – orange circle, male (M) – blue square, combined (F+M) – gray diamond, vehicle – open, CGRP – closed. B, OVAR significantly increased anxiety in all mice treated with IP vehicle (p = 0.002). B, OVAR significantly increased anxiety indexes in all mice treated with IP vehicle, and anxiety indexes also increased from vehicle to CGRP. OVAR further increased anxiety in CGRP-treated mice. **C**, Changes in anxiety indexes strongly correlated with time spent in the closed arms. Females were more sensitive to the effects of OVAR and CGRP than males. **D**, OVAR-induced anxiety strongly affected the number of center bouts detected in females and males during IP vehicle testing, and IP CGRP affected center bouts in females even without OVAR. P-values and asterisks are listed: ^*^p ≤ 0.05, ^**^p ≤ 0.01, ^***^p ≤ 0.001, ^****^p ≤ 0.0001.

### Rotarod

Dynamic balance was assessed with the Rotarod (Columbus Instruments) configured with either a mouse dowel (r = 1.5 cm) or a rat dowel (r = 3.5 cm). Mice were tasked to maintain balance and gait on the different dowels rotating from 5 to 44 rpm at an acceleration step of 2.4 rpm every 4 seconds. Latency to fall (LTF) is measured when mice fall from the dowel. Three days of rotarod testing were performed. The first day was a training day where mice were tested for 6 to 8 trials. On the second day, mice were briefly trained and were then injected with vehicle control. Twenty minutes after the injection, mice were tested for 3 trials (pre-VC) and were then stimulated with OVAR for 30 seconds. Mice were then immediately tested for 3 trials on the rotarod after the challenge (post-VC). Approximately 10-30 seconds pass in between subsequent trials during pre-VC and post-VC tests. On day 3, the same mice were re-tested with the same methods but were instead injected with CGRP (**Fig. 2A**).

**Figure 2:**
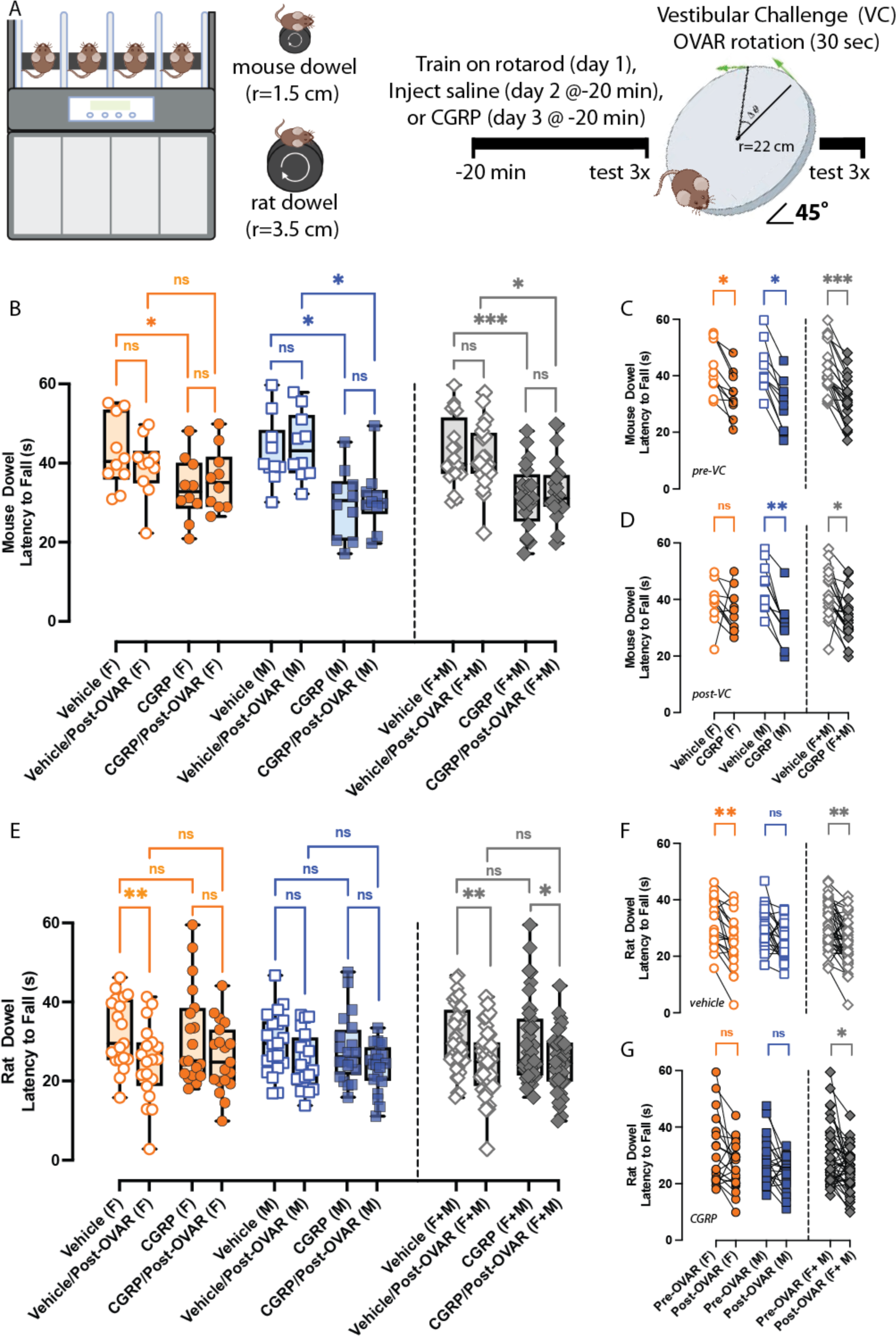
Calcitonin-Gene Related peptide (CGRP) and off-vertical axis rotation (OVAR) on rotarod configured with two different dowels. A, Rotarod configured with a mouse or rat dowel is shown, and an experimental timeline highlighting training and experiment days is provided. Twenty mice (10M/10F) were tested with the mouse dowel, and 39 mice (20M/19F) were tested with the rat dowel. Two-way repeated measures ANOVAs with Tukey multiple comparisons test were used to evaluate maximum latency to fall (s). **B-D**, Before the OVAR stimulus (when sexes are pooled), rotarod performance was reduced from IP vehicle to IP CGRP, and this change was also observed when comparing OVAR outcomes. A disrupted pre-VC rotarod ability due to CGRP is observed in both sexes. After the OVAR, only males indicated a significant difference between IP vehicle and IP CGRP. **E-G**, Opposite of mouse dowel studies, rat dowel studies indicated an effect of OVAR but not systemic CGRP. OVAR reduced rotarod ability in mice treated with IP vehicle and when treated with IP CGRP when sexes were pooled. In female mice specifically, max LTFs dropped from vehicle to vehicle/post-OVAR, but this was not seen in males. P-values and asterisks are listed: ^*^ p ≤ 0.05, ^**^p ≤ 0.01, ^***^p ≤ 0.001, ^****^p ≤ 0.0001.

### Vestibular Challenge – Off-Vertical Axis Rotation (OVAR)

In this study, a two-cage rotator (cage dimensions: 12.7 cm x 8.9 cm) was built to impose off-vertical axis rotation (60 rpm, 45° tilt from the vertical for 30 seconds) as a vestibular stimulus to mice during elevated plus maze and rotarod experiments. The mice are secured 22 cm from the rotational axis, and the device can rotate two mice at a time.

### Data Analysis and Statistics

Elevated plus maze data were analyzed by assessing time spent in closed arms as a percentage, the number of bouts through the center platform, and a calculated anxiety index (equation found in **Fig. 1A**). For rotarod assessment, the max latency to fall (LTF) of the three trials per testing condition was determined per mouse and indicated the mouse’s best effort for that testing condition. A group average of the max LTFs and standard error (mean + SEM) was computed for comparisons. Statistical analyses were performed using GraphPad Prism 9.5. Two-way repeated measures analysis of variance (RM-ANOVA) were primarily computed for females or males across elevated plus maze and rotarod testing, with the following factors: IP CGRP vs IP vehicle, and before versus after the vestibular challenge. Tukey multiple comparisons test was the preferred post hoc analysis for RM-ANOVAs. Unpaired t-tests were only used when comparing differences in rotarod ability due to dowel lengths after IP vehicle treatment. Statistical significance was set at p ≤ 0.05 for all analyses. F-values and p-statistics for RM-ANOVAs across behavioral outcomes can be found In **Table 1**.

**Table 1:** Two-way repeated measure ANOVAs were computed with Tukey multiple comparisons test as the post hoc analysis (Tukey adjusted p-values not shown). F values and associated p-values are provided across the behavioral outcomes assessed in this study: anxiety index (au), time in closed arms (%), bouts through center arm (au), rotarod configured with mouse dowel (s), and rotarod configured with rat dowel (s). ANOVAs were computed in females, males, and when sexes were pooled (F+M).

| Biological Sex | Behavioral Outcome | F (DF<sub>n</sub>, DF<sub>d</sub>) | P-val |
| --- | --- | --- | --- |
| Female | Anxiety Index | CGRP: $F(1.00, 7.00) = 67.11$,<br>OVAR: $F(1.00, 7.00) = 53.97$, | $p < 0.001$<br>$p < 0.001$ |
| | Time in Closed Arm (%) | CGRP: $F(1.00, 7.00) = 31.59$,<br>OVAR: $F(1.00, 7.00) = 129.1$, | $p < 0.001$<br>$p < 0.001$ |
| | Bouts Through Center Arm | CGRP: $F(1.00, 7.00) = 14.88$,<br>OVAR: $F(1.00, 7.00) = 380.8$, | $p = 0.006$<br>$p < 0.001$ |
| | Rotarod- mouse dowel | CGRP: $F(1.00, 9.00) = 9.71$,<br>OVAR: $F(1.00, 9.00) = 0.12$, | $p = 0.01$<br>$p = 0.73$ |
| | Rotarod - rat dowel | CGRP: $F(1.00, 18.00) = 0.02$<br>OVAR: $F(1.00, 18.00) = 14.34$ | $p = 0.9$<br>$p = 0.001$ |
| Male | Anxiety Index | CGRP: $F(1.00, 7.00) = 2.77$,<br>OVAR: $F(1.00, 7.00) = 30.09$, | $p = 0.14$<br>$p < 0.001$ |
| | Time in Closed Arm (%) | CGRP: $F(1.00, 7.00) = 16.38$,<br>OVAR: $F(1.00, 7.00) = 76.02$, | $p = 0.005$<br>$p < 0.001$ |
| | Bouts Through Center Arm | CGRP: $F(1.00, 7.00) = 7.63$,<br>OVAR: $F(1.00, 7.00) = 135.3$, | $p = 0.03$<br>$p < 0.001$ |
| | Rotarod- mouse dowel | CGRP: $F(1.00, 9.00) = 19.40$,<br>OVAR: $F(1.00, 9.00) = 1.74$, | $p = 0.002$<br>$p = 0.22$ |
| | Rotarod - rat dowel | CGRP: $F(1.00, 19.00) = 1.21$<br>OVAR: $F(1.00, 19.00) = 7.13$ | $p = 0.29$<br>$p = 0.01$ |
| Female + Male | Anxiety Index | CGRP: $F(1.00, 15.00) = 21.38$,<br>OVAR: $F(1.00, 15.00) = 82.01$, | $p < 0.001$<br>$p < 0.001$ |
| | Time in Closed Arm (%) | CGRP: $F(1.00, 15.00) = 42.38$,<br>OVAR: $F(1.00, 15.00) = 202.8$, | $p < 0.001$<br>$p < 0.001$ |
| | Bouts Through Center Arm | CGRP: $F(1.00, 15.0) = 22.30$,<br>OVAR: $F(1.00, 15.00) = 431.0$, | $p < 0.001$<br>$p < 0.001$ |
| | Rotarod- mouse dowel | CGRP: $F(1.00, 19.00) = 25.20$,<br>OVAR: $F(1.00, 19.00) = 0.17$, | $p < 0.0001$<br>$p = 0.69$ |
| | Rotarod - rat dowel | CGRP: $F(1.00, 38.00) = 0.19$<br>OVAR: $F(1.00, 38.00) = 20.98$ | $p = 0.67$<br>$p < 0.0001$ |

## Results

### CGRP and OVAR’s impact on elevated plus maze activity

In sixteen (8M/8F) C57BL/6J mice, we evaluated differences in anxiety indexes (**Fig. 1B**), time spent in closed arms (**Fig. 1C**), and the number of bouts through the center (**Fig. 1D**) by examining the effects of IP CGRP and OVAR as the vestibular challenge (VC). OVAR significantly increased anxiety in all IP vehicle-treated mice (p = 0.002). Anxiety indexes also increased from IP vehicle to IP CGRP prior to the OVAR (p = 0.02), and the OVAR further increases anxiety in CGRP-treated mice (p = 0.003) (**Fig. 1B**). These changes in anxiety indexes strongly correlate with time spent in the closed arms (**Fig. 1C**). Notably, all mice - on average - spend ∼52% of their time in the closed arms, but this percentage increases to ∼70% due to IP CGRP, ∼78% due to OVAR (IP vehicle injection), and ∼90% when mice are treated with IP CGRP and OVAR. When examining if sex-specific differences were present in anxiety outcomes, we observed that OVAR caused an increase in anxiety indexes (p = 0.03) for females and a trending effect in males (p = 0.09) (**Fig. 1B**). OVAR-induced anxiety correlated with an increased percentage of time spent in the closed arms for females (p < 0.001) and males (p = 0.02). Before OVAR, IP CGRP caused significant increases in female anxiety indexes (p = 0.01) and correlated with increased time in the closed arms (p = 0.002) but these observations were not seen in males (**Fig. 1C**). Additionally, CGRP/Post-OVAR anxiety indexes were higher than only CGRP-induced anxiety in females (p = 0.02) and correlated with increased time spent in the closed arms (p = 0.007). While the analyses on anxiety indexes and time spent in closed arms were robust in females, we did observe CGRP-treated males to spend increased time in the closed arms after the OVAR (p = 0.04), though this observation did not translate to a significant increase in male anxiety indexes. Additionally, we quantified the number of bouts through the center platform as an indicator of movement, because each bout indicates one instance where a mouse moved from an open to a closed arm and vice versa. In **Fig. 1D**, when sexes were pooled, IP vehicle-treated mice experienced a 2.8x fold reduction in center bouts after OVAR stimulation (p < 0.001). Likewise, a 3.2x reduction in center bouts was observed after OVAR when mice were treated with IP CGRP (p < 0.001). More specifically in a particular sex, OVAR strong affected the number of center bouts detected in females (p < 0.001) and in males (p < 0.001) during IP vehicle testing. After the OVAR in IP CGRP-treated mice, females exhibited significantly reduced center bout activity (p = 0.009) and so did males (p < 0.001). Even without OVAR, IP CGRP affected center bouts in females (p = 0.01) and this was a trend seen in males (p = 0.06). These results show CGRP and a vestibular challenge can independently elicit anxiety-like behavior in mice with a female preponderance in observed outcomes, but the use of both stressors is additive and elicits even stronger anxiety in mice.

### Differences in C57B6/J’s rotarod ability tested on different dowel diameters

While OVAR Is used as the primary vestibular challenge, we wanted to observe how balance function differs when mice perform rotarod on different dowel lengths (1.5 cm vs 3.5 cm radius). We hypothesized that mice would be further challenged on the rat dowel, because to maintain balance on a larger diameter dowel rotating at the same parameters (in rpm) as a smaller dowel, a greater distance of the larger dowel’s circumference must be displaced per second. We calculated that the mouse had to travel ∼2.3x more distance on the rat dowel per second of rotation at the lowest (5 rpm) and the highest (44 rpm) speeds, ultimately becoming more of a challenge. This difficulty was reflected in the data. We observed IP vehicle-treated mice on the mouse dowel exhibited higher max LTFs (n = 20, 42.62 ± 1.93 s) compared to mice tested on the rat dowel (n = 39, 30.75 ± 1.33 s), highlighting a difference in best performance by -11.87 ± 2.31 seconds when naïve mice are challenged to do rotarod on a larger dowel diameter (unpaired t-test, t = 5.13, df = 57, p < 0.0001). Due to this disparity in best performance between dowel lengths, we attempted to calibrate the rat dowel so that instead of matching the mouse dowel’s rpm, it would match the displaced distance per second. We dedicated twenty naïve, age-matched mice (10M/10F) to this task, and observed mice to exhibit significantly higher max LTFs (greater than 90 seconds on average per trial) when rat dowel parameters were altered (data not shown). We surmised that mouse gait cannot be easily tuned by modulating the displaced distance of the dowel and that other factors must be at play (e.g., attention, position of paws in respect to curved dowel surface), and thus did not explore this calibration further.

### CGRP and OVAR’s effects on mouse dowel-rotarod

Twenty mice (10M/10F) were assessed for rotarod ability on the mouse dowel, before and after OVAR as the vestibular challenge (VC). In all mice prior to the OVAR stimulus (pre-OVAR), IP CGRP significantly reduced rotarod performance with a mean reduction of 11.01 ± 2.07 s (p = 0.0002) compared to IP vehicle (**Fig. 2B-D**). This impaired performance due to systemic CGRP was still observed after OVAR, as post-OVAR outcomes showed a decrease from IP vehicle to IP CGRP (p = 0.03). A disrupted pre-VC rotarod ability due to CGRP is observed in both sexes, with a mean reduction of 9.02 ± 2.47 s (p = 0.02) in females and of 13.00 ± 3.32 (p = 0.02) in males. When comparing post-OVAR outcomes, only males indicated a significant difference between IP vehicle and IP CGRP (p = 0.009).

### CGRP and OVAR’s effects on rat dowel-rotarod

Thirty-nine mice (20M/19F) were used to assess CGRP and OVAR’s effects on rotarod configured with a rat dowel (r = 3.5 cm). When sexes are pooled, IP CGRP had no effect but OVAR significantly affected performance in all mice (**Fig. 2E-G**). OVAR reduce rotarod ability in mice treated with IP vehicle by 5.92 ± 1.44 s (p = 0.001) and when treated with IP CGRP by 4.92 ± 1.72 s (p = 0.03). In female mice specifically, OVAR impacted performance after IP vehicle by 7.51 ± 1.87 s (p = 0.004). OVAR did not significantly impact male rotarod ability on the rat dowel after IP vehicle, nor were post-OVAR outcomes strongly different after IP CGRP testing.

## Discussion

Preclinical research in migraine – via rodent models – has provided evidence that biological sex can drive the occurrence of certain mouse surrogate behaviors that resemble clinically reported symptoms, known to arise more often in women than men. In this study through the elevated plus maze (EPM) test, we show that intraperitoneally delivered (IP) CGRP causes a majority of female mice to spend increased time spent in the closed arms and exhibit significantly higher anxiety indexes compared to their baseline, but this change in anxiety indexes was not significantly seen in males. Consistent with our results, a prior study indicated that central, intracerebroventricular (i.c.v.) administration of CGRP from 250 ng to 1 ug decreases open-field activity of adult male Wister rats in a dose-dependent manner (Kovács et al. 1999). A previous study on mutant mice, with an existing severe vestibular deficit showed increased anxiety when tested on the elevated plus maze and also were impaired on the mouse dowel rotarod. Interestingly, daily vestibular training improved both anxiety and rotarod behaviors (Shefer et al. 2015) in these mutant mice.

Motion-induced anxiety was more pronounced in female mice when were tested with the EPM after being stimulated with a30 second off-vertical axis rotation (OVAR) as a vestibular challenge. Understandably, both sexes experience a significant increase in motion-induced anxiety (a near 38% increase in anxiety-indexes) due to CGRP and OVAR when compared to their normal baseline anxiety - measured after vehicle administration (no vestibular challenge). We also observed reduced movement in the EPM when we measured a mouse’s bouts through the center, as mice treated with OVAR were not as active as their pre-OVAR state after IP vehicle or IP CGRP injections. This motion-induced anxiety and reduced activity observed during EPM is of clinical importance, as a clear, positive correlation between anxiety, depression, and vertigo is observed in patients with vestibular disorders (Omara et al. 2022), and so studying this link in mouse models provides insight into this correlation. The EPM is also a suitable assay as a surrogate behavior for agoraphobia in mouse models for migraine, as mice spend more time in the closed arms when they exhibit anxiety-like behaviors, and mirrors patients’ tendencies to avoid open spaces during vestibular migraine attacks (Kutay et al. 2017).

Our EPM data is consistent with previous reports in rodents that suggest a female preponderance towards anxiety. Experiments using the open field assay and unpredictable, sound stress highlighted a female likelihood to anxiety-like behaviors (Viero et al. 2022; Wang et al. 2022). Outside of anxiety but still related to the CGRP’s sexually dimorphic effects, administration of CGRP to the spinal cord (intra-thecal) caused a dose-dependent, longer-lasting hind paw allodynia in female rodents, as opposed to a more momentary effect in males (Paige et al. 2022).

The use of OVAR as a vestibular provocation is rarely used in rodent studies, and here we show that even a brief, vestibular stimulus can bring out significant anxiety and balance complications in the wildtype C57BL/6J strain.

Anxiety is a risk factor for the progression of episodic to chronic migraine, and elevated CGRP levels have been shown to alter neuronal activity that is linked to anxiety behaviors (Durham 2016). The results here further elucidate CGRP’s role in motion-induced anxiety and balance function in mouse models, and should be further explored in the presence of CGRP-targeted therapeutics. Peripherally administered CGRP-signaling antagonists have been widely observed to reduce CGRP-induced migraine-like sensitivities in rodents and humans, and therefore would be hypothesized to attenuate anxiety caused by IP CGRP (Araya et al. 2019; Raffaelli, Neeb, and Reuter 2019).

To conclude, our study extends our current understanding of CGRP-induced anxiety by incorporating a motion sensitivity component, and uniquely evaluates mouse balance function to a degree not observed in prior rotarod studies. We observed a female preponderance in both CGRP and motion-induced anxiety-like behaviors, suggesting that the role of CGRP in migraine’s anxiety symptoms can be recapitulated in preclinical models. The results of this study provide support to claims that CGRP has a pertinent role in motion-induced anxiety and dynamic imbalance, and warrants the potential use of anti-CGRP therapies for the treatment of these symptoms.

## Acknowledgments

We would like to acknowledge Travis Kovitz, Katharine Bachmann, and the Behavioral Sciences Facility Core at the University of Rochester for data collection of elevated plus maze experiments. Figures created with BioRender.com. This work was fully supported by NIH R01DC017261.

